## Supplemental Materials & Methods for "Iroki: automatic customization and visualization of phylogenetic trees"

Supplementary Materials & Methods

#### Contents

**1 Bacteriophage proteomes, taxonomy, and host phyla** **B–1**
1.1 Collecting phage genomes . . . . . B–1
1.2 Building proteomic tree . . . . . B–1

**2 Bacterial community diversity and prevalence of *E. coli* in beef cattle** **B–1**
2.1 Generating the tree . . . . . B–1
2.2 Generating the mapping file . . . . . B–2

**3 *Tara* Oceans viromes** **B–2**
3.1 Collecting RNR sequences . . . . . B–2
3.2 Building the count table and trees . . . . . B–2
3.3 Calculating sample covariate correlation . . . . . B–3
3.4 PCA biplot . . . . . B–3

**4 Canvas viewer for large trees** **B–3**

**5 Supplementary Figures** **B–4**

This document contains supplementary materials and methods for the manuscript *Iroki: automatic customization and visualization of phylogenetic trees*.

### 1 Bacteriophage proteomes, taxonomy, and host phyla

#### 1.1 Collecting phage genomes

Phage genomes were collected from the Virus-Host DB last updated on 2018-08-08 [1]. This version of the database pulled from RefSeq [2] release 90 (September 10, 2018) and GenBank [3] release 227.0 (August 15, 2018). Viral genomes were selected that matched the following criteria: (1) viral genome length is greater than 20,000 bases, and (2) the host phyla is one of Actinobacteria (688 viruses with this host phylum), Bacteroidetes (39 viruses), Cyanobacteria (98 viruses), Firmicutes (591 viruses), or Proteobacteria (1038 viruses). Viral genomes matching this criteria yielded viruses from only a few families: Ackermannviridae (21 viruses in this family), Myoviridae (639 viruses), Podoviridae (421 viruses), and Siphoviridae (1373 viruses). In total, 2,454 viral genomes were selected. Scripts used in this process can be found on Zenodo (`https://doi.org/10.5281/zenodo.3458510`).

#### 1.2 Building proteomic tree

The proteomic tree was built using ViPTree, a software package that automates proteomic tree construction [4]. ViPTree first uses tBLASTx to generate normalized similarity scores between all genomes ($S_G$; $0 \leq S_g \leq 1$). Then, to build the tree, BIONJ clustering is performed on genomic distances ($1 - S_G$).

### 2 Bacterial community diversity and prevalence of *E. coli* in beef cattle

#### 2.1 Generating the tree

A subset of operational taxonomic units (OTUs) were selected from a previous study examining the diversity of the bacterial community associated with beef cattle hide [5]. A Mann-Whitney U test comparing OTU abundance between STEC positive and STEC negative samples was performed using QIIME's `group_significance.py` script (MacQIIME version 1.9.1, `http://www.wernerlab.org/software/macqiime`) [6]. Cluster representative sequences from any OTU with a $p$-value $< 0.2$ were selected. These sequences were aligned with the SILVA ACT service with the default settings against the SILVA SSU database (`https://www.arb-silva.de/aligner/`) [7]. SILVA ACT was also used to generate an approximate-maximum likelihood tree with FastTree (GTR model, Gamma rate model for likelihoods), and to predict OTU taxonomy.

#### 2.2 Generating the mapping file

The mapping file specifies the heights and color of two series of bars that are based on OTU abundance data. For the first bar series (Fig. 2, inner ring of purple bars), bar height is based on $log + 1$ transformation of overall OTU abundance. Each OTU was ranked based on this abundance, and these ranks were used to generate a purple color palette (low abundance OTUs: light purple, high abundance OTUs: dark purple) using Iroki's color gradient generator (`https://www.iroki.net/`).

The second bar series (Fig. 2, outer ring of brown/blue bars) shows the abundance ratio of the OTUs between STEC negative and STEC positive samples. Let $A_i^-$ be the mean abundance of OTU $i$ in STEC negative samples, $A_i^+$ be the mean abundance of an OTU $i$ in STEC positive samples, and $R = A_i^- / A_i^+$, be the abundance ratio of OTU $i$ in STEC negative to STEC positive samples. Then the bar height is calculated as follows:

$$H = \begin{cases} R - 1, & \text{if } R \geq 1 \\ 1 - (1/R), & \text{otherwise} \end{cases} \tag{1}$$

In this way, OTUs that have approximately the same abundance in STEC positive and negative samples will have a height of about zero, OTUs with a higher abundance in samples without STEC will have high magnitude positive heights (brown bars facing outwards towards the leaf labels), and OTUs with a higher abundance in samples with STEC will have high magnitude negative heights (blue bars facing inwards towards the tree root).

### 3 *Tara* Oceans viromes

#### 3.1 Collecting RNR sequences

RNR sequences were identified in 44 *Tara* Oceans viromes via homology search using MMSeqs2 [8] (`mmseqs` commit: `2cfdedc95f6a998826f45a7594971751a5e535f3`). Three viromes containing less than 50 RNR sequences were not used in further analysis. In total, 5,470 RNR sequences from 41 samples were collected.

#### 3.2 Building the count table and trees

A presence/absence count table (biom table) was constructed according to the following rule: an RNR sequence $r$ was considered present in a sample $s$ if sequence $r$ originated from sample $s$. Next, RNR sequences were aligned with the MAFFT [9] v7.388 plugin (default settings) in Geneious R10. The resulting alignment was manually inspected to ensure key conserved residues were properly aligned. An approximate-maximum likelihood tree was inferred from this multiple sequence alignment using the FastTree v2.1.10 with double precision (default settings) [10]. This tree was then used in conjunction with the count table to calculate unweighted UniFrac distance [11] between samples using QIIME (MacQIIME v1.9.1) [6]. Calculating distance in this way generates distance between samples based solely on phylogenetic relatedness of the samples' RNR proteins.

Finally, average-linkage hierarchical clustering was performed on the samples using the unweighted UniFrac distance matrix in R [12] using the `hclust` function. The R package `ape` [13] was used to export tree to Newick format.

##### 3.3 Calculating sample covariate correlation

To determine which sample covariates were significantly correlated with viral community structure, distance matrices were constructed from available metadata using QIIME's `distance_matrix_from_mapping.py` script. Then, the count table was rarefied to 50 sequences to mitigate the affect of sequencing depth on correlation calculations, and this table was used to calculate unweighted UniFrac distance between samples. Finally, Mantel tests were performed with QIIME's `compare_distance_matrices.py` script using this UniFrac distance matrix in conjunction with the sample covariate distance matrices based on environmental factors.

##### 3.4 PCA biplot

A PCA biplot was made to show how *Tara* Ocean viromes related to one another with respect to the environmental parameters included in Fig. 3. PCA of samples included in Fig. 3 was calculated and plotted using the R package `biplotR` (code repository: https://github.com/mooreryan/biplotr; archived in Zenodo: [14]). Data were centered and scaled prior to PCA. Samples outside of clusters A, B, and C were excluded from Fig. 4 in the main text. The full biplot is shown in Supplementary Fig. S1.

#### 4 Canvas viewer for large trees

Given the substantial growth in sequence databases and the burgeoning capacity of DNA sequencers, it is now possible to obtain millions of sequences and thousands of samples for analysis. Such datasets often generate large trees; however, rendering such large trees without collapsing nodes is challenging. To address this, Iroki employs an HTML5 Canvas straight-to-png tree viewer with the ability to display trees with millions of leaf nodes (test tree was generated with the `rtree` function of the `ape` R package [13]) in under 20 seconds on our test machine (MacBook Pro, 2.66 GHz Intel Core i7, 8GB RAM, with 64bit Google Chrome web browser version 76.0.3809.132) (Supplementary Fig. S2). While the Canvas tree viewer currently lacks many of the features of the Iroki's SVG viewer, the ability to render such huge trees may still prove valuable.

To illustrate the utility of this feature, the GreenGenes [15] ”two-study filter” tree (331,550 sequences) was obtained from the FastTree website for visualization using Iroki (Supplementary Fig. S3). Briefly, full length SSU rRNA sequences from GreenGenes were collected and clustered based on divergence from an ancestor in a minimum-evolution tree. Sequences were kept only if they were in a cluster with sequences from two different studies or if they were named isolates. Sequences were then aligned with NAST and the phylogeny inferred with FastTree [10] (see www.microbesonline.org/fasttree for full details).

### 5 Supplementary Figures

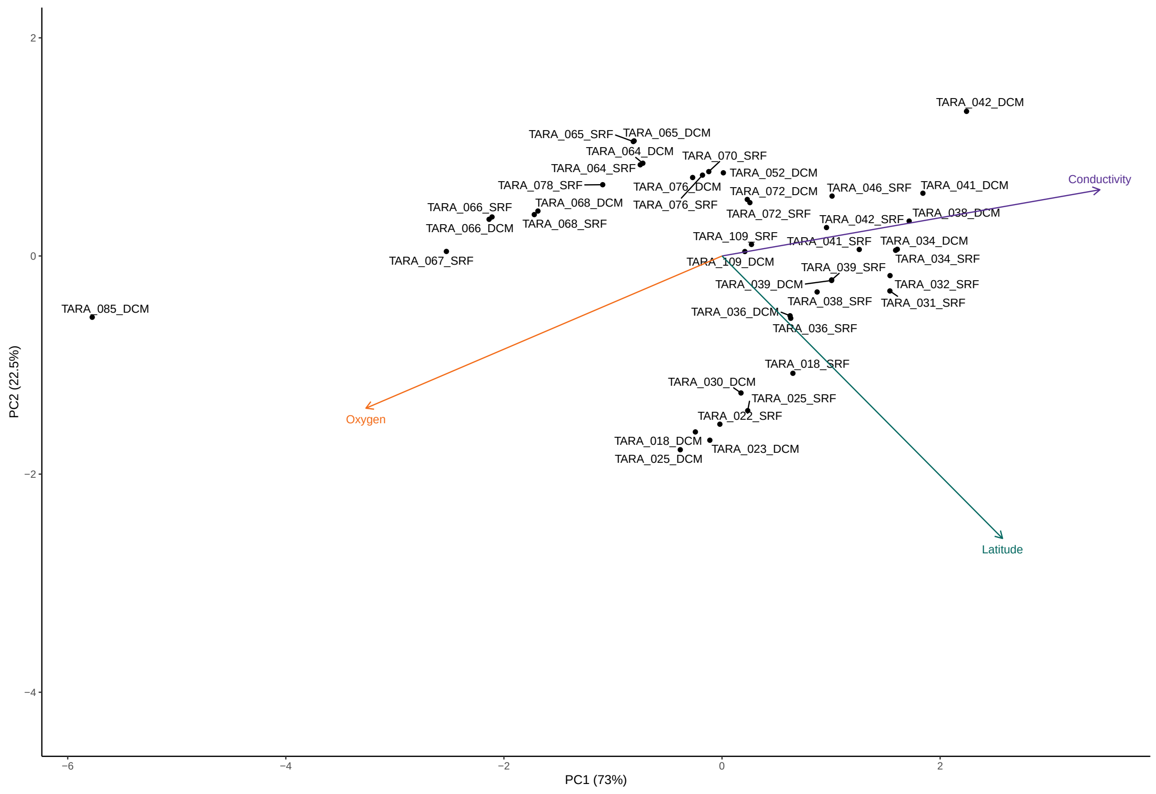


**Figure S1: Full PCA biplot of *Tara* Oceans viromes.** Principal components analysis biplot of 41 *Tara* Oceans viromes based on sample oxygen, conductivity, and latitude.

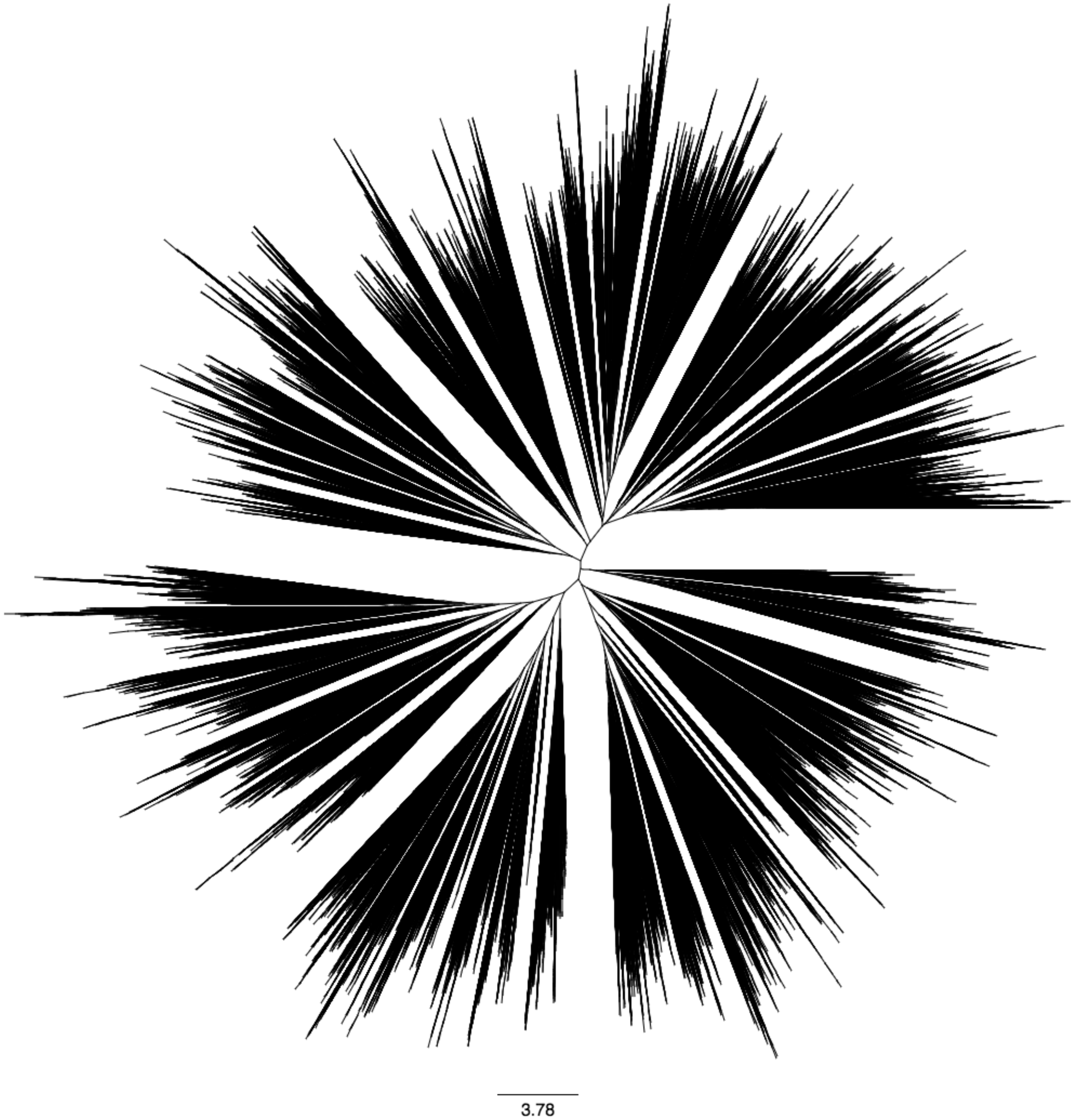


**Figure S2: Random branch length tree**. A 1,000,000 leaf tree with random branch lengths generated with `rtree` (using `runif` with default arguments for branch lengths) from the ape R package. Tree was rendered with Iroki's Canvas tree viewer.

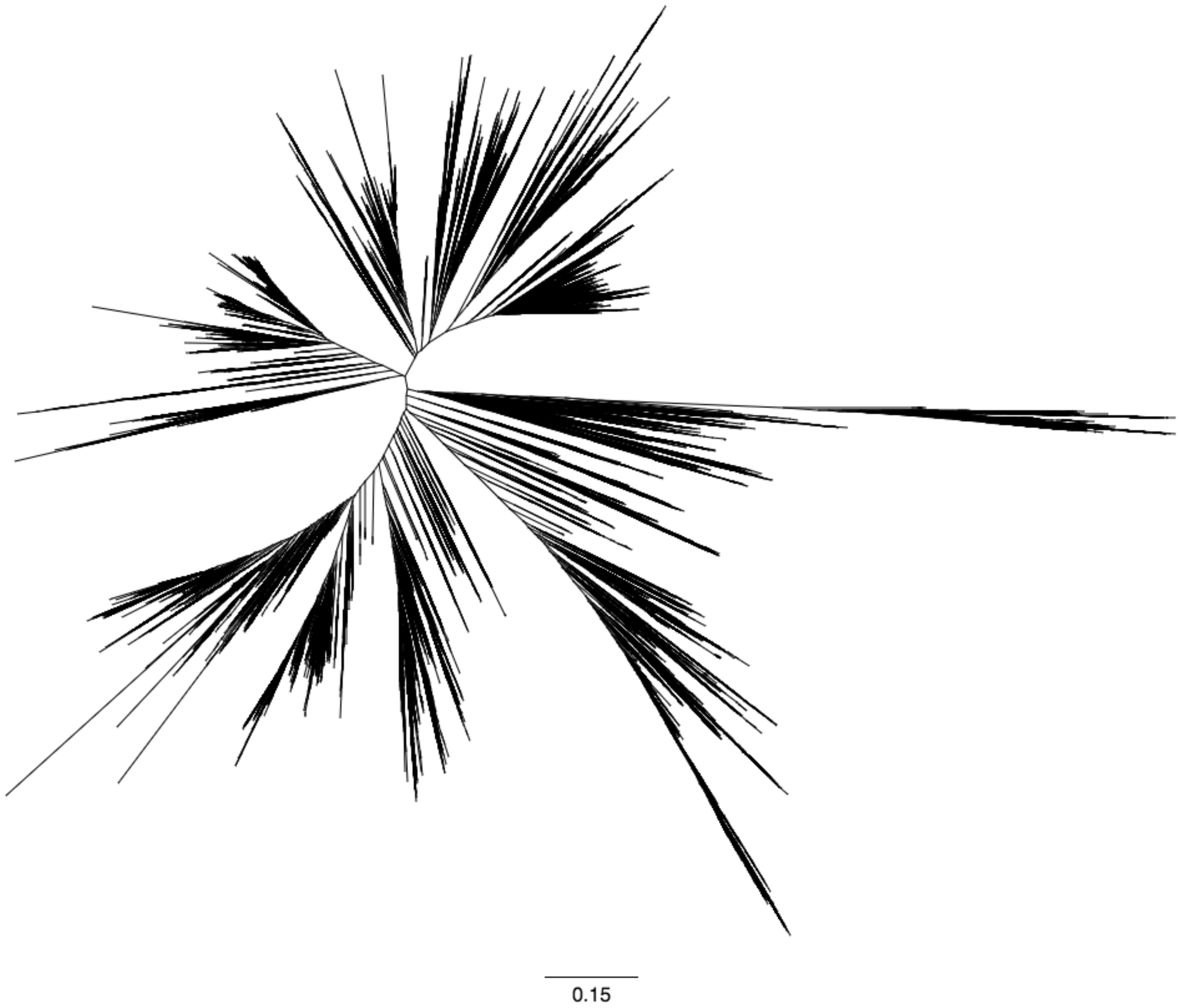


**Figure S3: GreenGenes SSU rRNA tree.** A collection of 331,550 full length SSU rRNA sequences from GreenGenes rendered with Iroki's Canvas tree viewer.

[6] J Gregory Caporaso, Justin Kuczynski, Jesse Stombaugh, Kyle Bittinger, Frederic D Bushman, Elizabeth K Costello, Noah Fierer, Antonio Gonzalez Peña, Julia K Goodrich, Jeffrey I Gordon, Gavin A Huttley, Scott T Kelley, Dan Knights, Jeremy E Koenig, Ruth E Ley, Catherine A Lozupone, Daniel McDonald, Brian D Muegge, Meg Pirrung, Jens Reeder, Joel R Sevinsky, Peter J Turnbaugh, William A

Walters, Jeremy Widmann, Tanya Yatsunenko, Jesse Zaneveld, and Rob Knight. QIIME allows analysis of high-throughput community sequencing data. *Nature Methods*, 7(5):335–336, May 2010. ISSN 1548-7091. doi: 10.1038/nmeth.f.303. URL http://www.nature.com/doifinder/10.1038/nmeth.f.303.
